# The effect of non-linear competitive interactions on quantifying niche and fitness differences

**DOI:** 10.1101/2021.08.30.458252

**Authors:** Jurg Spaak, Remi Millet, Po-Ju Ke, Andrew D. Letten, Frederik De Laender

## Abstract

The niche and fitness differences of modern coexistence theory separate mechanisms into stabilizing and equalizing components. Although this decomposition can help us predict and understand species coexistence, the extent to which mechanistic inference is sensitive to the method used to partition niche and fitness differences remains unclear. We apply two alternative methods to assess niche and fitness differences to four well known community models. We show that because standard methods based on linear approximations do not capture the full community dynamics, they can sometimes lead to incorrect predictions of coexistence and misleading interpretations of stabilizing and equalizing mechanisms. Conversely, a more recently developed method to decompose niche and fitness differences, that accounts for the full nonlinear dynamics of competition, consistently identifies the correct contribution of stabilizing and equalizing components. This approach further reveals that when the true complexity of the system is taken into account, essentially all mechanisms comprise both stabilizing and equalizing components. Amidst growing interest in the role of non-additive and higher-order interactions in regulating species coexistence, we propose that the effective decomposition of niche and fitness differences will become increasingly reliant on methods that account for the inherent non-linearity of community dynamics.

## Introduction

Niche and fitness differences are two central concepts in coexistence theory which help us to understand species coexistence. Whether or not species coexist depends on how much species limit each other’s growth compared to their own growth. From a mechanistic standpoint, various factors such as resources (Letten *et al*., 2017), predators Chesson & Kuang (2008) or mutualists (Johnson, 2021), can limit population growth. Population sizes will in turn influence these factors (Meszéna *et al*., 2006) (e.g. by depleting resources or by boosting the population size of their predators), as such creating the regulating feedback underpinning growth limitation. Thus, mechanistically, niche differences (1-niche overlap) measure how independent the feedback loops of two species are. If the feedback loops are completely independent, then niche differences are 1, conversely, if all feedback loops, intra and interspecific, are equivalent niche differences are 0. Fitness differences measure the relative strength of the feedback loops. Species will coexist when niche differences overcome fitness differences (Adler *et al*., 2007; Chesson, 2013).

Unfortunately, computing niche and fitness differences taking into account the full complexity of limiting factors is not a straightforward task. Some available methods therefore apply to simple models only (Spaak & De Laender, 2020), where the details of population regulation are omitted. One often applied method is to reformulate the equilibrium conditions of a community model in linear terms, equivalent to to the equilibrium conditions of a linear Lotka-Volterra model (Letten *et al*., 2017; Godoy *et al*., 2014; Johnson, 2021). This writes the equilibrium condition as a linear function of the competitor’s densities, which then act as limiting factors, in lieu of the actual factors that underpin species interactions. Often, this includes various assumptions on the time-scales at which the limiting factors vary, as well as on the persistence of these factors (Letten *et al*., 2017; Johnson, 2021). While these simplifying assumptions will by design be violated in certain conditions, we do not know the implications of such violations for our capacity to understand coexistence. Such knowledge is important for application of coexistence theory, as is choosing the appropriate definition of niche and fitness difference (Godwin *et al*., 2020).

Here, we examine to what extent including non-linear characteristics of community models increases our understanding of coexistence. To this end, we assemble four community models, three of which explicitly model limiting factors, and one which considers direct non-linear effects of population density on per-capita growth. We rewrite the equilibrium conditions of all four models with linear functions and compare how the resulting niche and fitness differences differ from those of the full models including limiting factors and direct non-linear interactions. Based on this comparison, we first develop criteria to test whether the approximated models affect our ability to understand coexistence. Then, we graphically represent the assumptions made by the approximation method and show that these simplifications can result in large deviations in community dynamics, especially away from equilibrium. Importantly, we show that linear approximations of non-linear dynamics can sometimes result in the misunderstanding of community dynamics, e.g. negative frequency dependence interpreted as positive frequency dependence and vice-versa. Finally, we also show that accounting for the full non-linear dynamics of competition can yield new insights into the effects of mortality and resource availability on niche and fitness differences.

## Methods

Niche and fitness differences of four community models have so far been computed by analyzing the linear equilibrium conditions: Species competing for substitutional resources with a Holling type 1 response (Chesson, 1990) and a Holling type 2 response (Letten *et al*., 2017), species competing for essential resources with a Holling type 2 response (Letten *et al*., 2017) and the annual plant model (Godoy & Levine, 2014), see table 1. For each of these models we used the niche and fitness differences (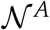 and 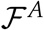) from the literature (Chesson, 1990, 2013, 2018). The superscript A denotes their derivation from **a**pproximated models. To compute niche and fitness differences of the full model (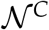 and 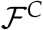) we use the method outlined in Spaak & De Laender (2020). The superscript *C* denotes their derivation from the full, non-linearised (**c**omplex) model. We will use 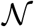 and 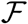 to denote niche and fitness differences based on any mathematical definition. All simulations and computations were done in Python 3.7.1 with NumPy version 1.15.4.

**Table 1:**
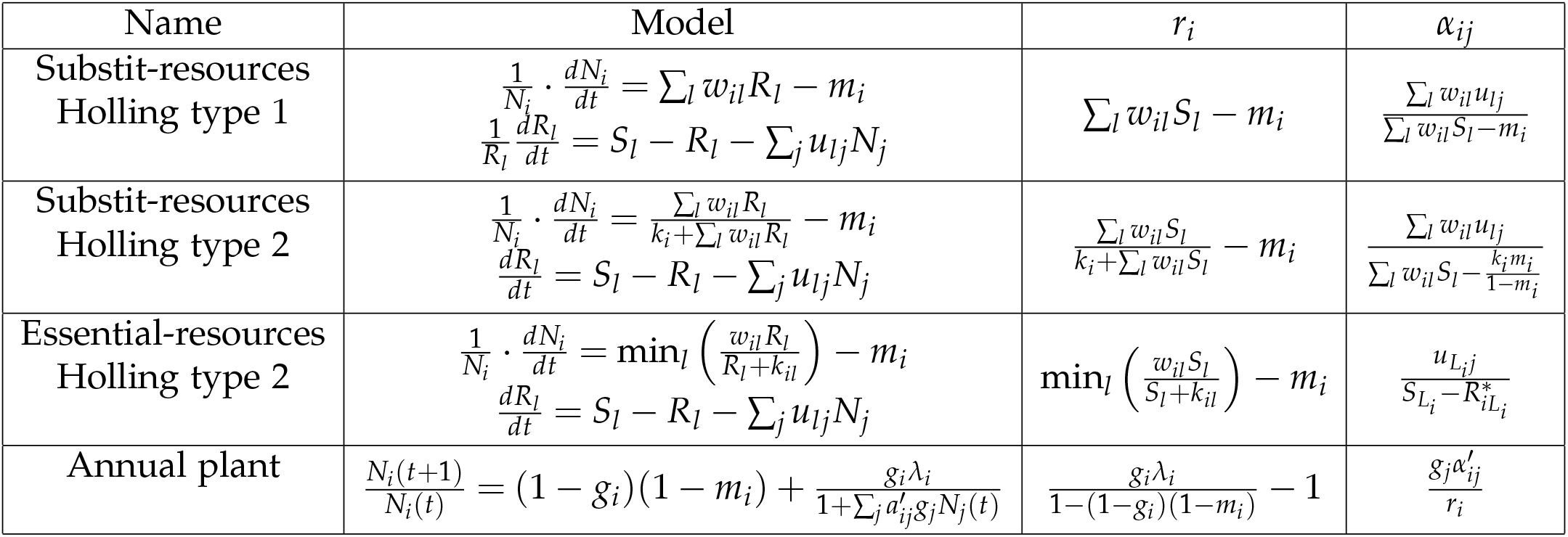
We consider four community models of which the full equations are given (‘Model’), as well as their linear representation of the equilibrium dynamics as reported before (Chesson, 2013; Godoy *et al*., 2014; Letten *et al*., 2017). The linearized versions of these models are reconstructed by replacing *r_i_* and *α_ij_* in equation 1 by the expressions in the corresponding columns. The first three models are resource explicit models, in the main text we focus on two species communities competing for two resources. The resource growth rates may be either in per-capita (as in the first model) or in total (as in the second and third model), this does not qualitatively change our results. We chose resource growth rates as used in Chesson (2013) and Letten *et al*. (2017). For these models *w_il_* is the conversion of resource to biomass, *u_li_* is the utilisation of resource *l* by species *i*, *m_i_* is the mortality rate, *S_l_* is the resource supply and *k_i_* respectively *k_il_* are half-saturation constants. Consistently, subscripts *i* and *l* stand for species and resources, respectively. Subscript *L_i_* is the index of the more limiting resource of species *i* in monoculture. In the annual plant model *g_i_* is the germination rate, *m_i_* is the seed mortality rate (traditionally the model uses the survival rate *S_i_* = 1 — *m_i_*), *λ_i_* is the net-production rate and 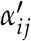 is the intra- or interspecific interaction.

**Table 2:** How 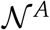 and 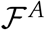 or 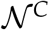 and 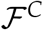 interpret coexistence across the four investigated models (columns). ✓indicates correct prediction respectively explanation, ✕ indicates wrong prediction or explanation. The annual plant model does not explicitly consider limiting factors, so there is no clear criteria for asserting that both species occupy the same niche, the two methods however agree perfectly, i.e. 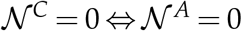, which we indicate with a “=” (see Appendix S5). Similarly, there is no clear criteria for asserting that a mechanism is stabilizing, and we found that the two definitions do not agree on this (indicated with “≠”).

| Criteria | Substit-resources<br>Holling type 1 |  | Substit-resources<br>Holling type 2 |  | Essential-resources<br>Holling type 2 |  | Annual plant |  |
| --- | --- | --- | --- | --- | --- | --- | --- | --- |
| | $\mathcal{N}^C, \mathcal{F}^C$ | $\mathcal{N}^A, \mathcal{F}^A$ | $\mathcal{N}^C, \mathcal{F}^C$ | $\mathcal{N}^A, \mathcal{F}^A$ | $\mathcal{N}^C, \mathcal{F}^C$ | $\mathcal{N}^A, \mathcal{F}^A$ | $\mathcal{N}^C, \mathcal{F}^C$ | $\mathcal{N}^A, \mathcal{F}^A$ |
| No interaction | ✓ | ✓ | ✓ | ✓ | ✓ | ✓ | ✓ | ✓ |
| Same niche | ✓ | X | ✓ | X | ✓ | X | = | = |
| Stabilizing mechanism | ✓ | X | ✓ | X | ✓ | X | ≠ | ≠ |

### Niche and fitness differences based on approximated models

The computation of the approximated niche and fitness differences 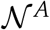 and 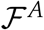 is based on rearranging the equilibrium condition of a community model, given by 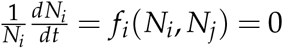, into a form that is equivalent to the equilibrium condition of a Lotka-Volterra community model, given by:

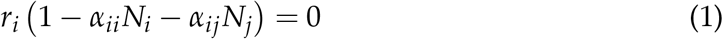

Where *N_i_* is the density of species *i*, *r_i_* is the intrinsic growth rate, *α_ii_* and *α_ij_* are the intra- and interspecific competition coefficients. This involves writing the *α_ij_* of equation 1 as a function of the model parameters. Generally speaking, the linear approximations are chosen to match the zero net growth isoclines (ZNGI), i.e. *f_i_* (*N_i_, N_j_*) = 0 ⇔ 1 – *α_ii_N_i_* – *α_ij_N_j_* = 0. However, the ZNGI may not be linear, in which case such a linear representation of the ZNGI will necessarily be an approximation. We show below, that the approximations do sometimes incorrectly approximate the equilibrium densities. The *r_i_* are equivalent to the intrinsic growth rates, i.e. *r_i_* = *f_i_* (0,0). Table 1 summarizes the linear representation for all these models. Given the interaction coefficients *α_ij_* we can compute 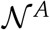 and 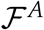 as (Chesson, 2018):

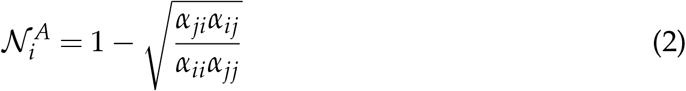

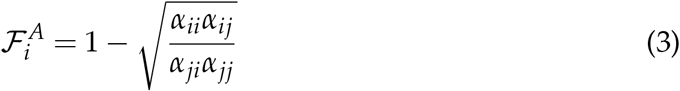

Note that we slightly change the definition of 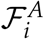 by defining 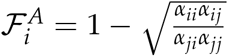 instead of the more usual 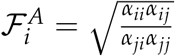. This change is purely aesthetic to ensure consistency between 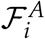 and 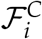, it does not affect the properties of 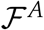.

### Niche and fitness differences based on the full model

We computed 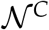 and 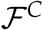 for the full model based on a recently developed method (Spaak & De Laender, 2020). Briefly, this method ensures that species with independent feedback loops have 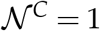, while species with equivalent feedback loops have 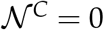. Thus, the method critically depends on whether invasion analysis can predict coexistence for the model it is applied to (Spaak & De Laender, 2020; Barabás *et al*., 2018; Chesson, 1994; Pande *et al*., 2019). The invasion growth rate of a species *i* is its growth rate when the resident species *j* is at its monoculture equilibrium density. More precisely, for a model given by the per-capita growth rate 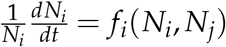 Spaak & De Laender (2020) define 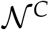 and 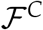 as

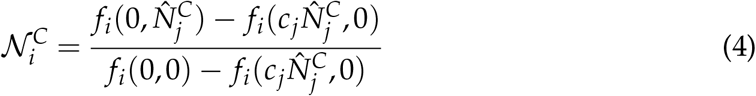

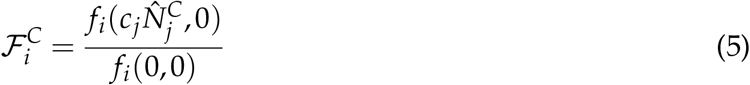

where 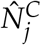 is, as above, the monoculture equilibrium density of species *j*. *c_j_* is a conversion factor that converts densities of species *j* to densities of species *i*, it ensures that both species have the same dependence on limiting factors, i.e. equally strong feedback loops. *c_j_* is the solution of the equation 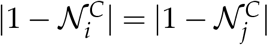

Equation 4 compares the actual invasion growth rate of species 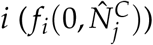 to the hypothetical invasion growth rate when the two species had independent feedback loops (*f_i_*(0,0)) and to another hypothetical invasion growth rate when the two species had identical feedback loops 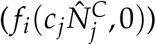.

### Predicting and understanding coexistence

For 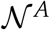 and 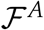 or, 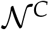 and 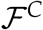 to predict coexistence they need to correctly predict the outcome of species interactions. For a two species community there are three such outcomes (Tilman, 1982).

A first outcome is priority effects, where the outcome of competition depends on the starting conditions, this is only possible with positive frequency dependence, i.e. negative niche differences (Ke & Letten, 2018). A second outcome is coexistence, where both species persist indefinitely, this is only possible with negative frequency dependence, i.e. positive niche differences. A third outcome is competitive exclusion, where the competitive dominant species excludes the other species. This is consistent with both negative or positive niche differences, and depends on the fitness differences 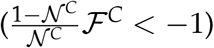. When 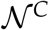 and 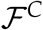 or, 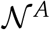 and 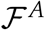 predict these three outcomes for a given model, we consider these to correctly *predict* coexistence.

For 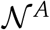 and 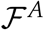 or, 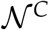 and 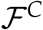 to *understand* coexistence, they need to reflect the mechanisms driving the outcome of species interactions. 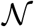 measure the difference between the niches of two species. When the two species do not interact with each other, because they occupy completely different niches, then 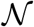 should reflect total niche differences, i.e. 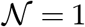 (Spaak & De Laender, 2020). Conversely, if two species occupy the same niche, then 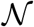 should be 0. Testing whether two species occupy the same niche is difficult in general, as one may not know whether the species do differ in any not-investigate niche axis. However, in the special cases where there is only one limiting factor, e.g. because species compete only for one resource or resources go extinct, species must occupy the same niche.

In addition, to *understand* species coexistence, 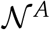 and 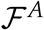 or, 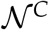 and 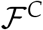 need to categorize a change in a model’s parameter as stabilizing and/or equalizing mechanism. That is, changes that increase niche differences are called stabilizing, while they are equalizing when they decrease fitness differences. For simplicity we will use the terms stabilizing and equalizing for both, increases and decreases in niche differences or fitness differences, respectively. Importantly, mechanisms can be both stabilizing and equalizing. We will call any mechanism that only affects niche or fitness differences purely stabilizing or equalizing, respectively. We tested for all model parameters whether they are stabilizing or equalizing according to 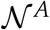 and 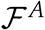 or, 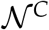 and 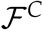.

## Results

Implicitly, 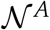 and 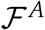 are linear approximations of the per-capita growth rates, i.e. *f_i_*(*N_i_*,*N_j_*) ≈ *r_i_* · (1 — ∑_*j*_*α_ij_N_j_*) (Fig. 1). Through this approximation we often loose in-formation about community dynamics, most notably higher order interactions (Grilli *et al*., 2017; Levine *et al*., 2017) or explicit interactions with limiting resources (Letten & Stouffer, 2019). Specifically, the linearisation does not account for substitutable resources to go extinct (Chesson, 1990; Letten *et al*., 2017), or for the fact that the identity of the essential resource that is limiting to a species can change during competition, independent of changes in the resource supply rates (Letten *et al*., 2017). As is to be expected, the linearisation sometimes gives a poor approximation of the growth rates, and of the invasion growth rate especially, for all four models (Fig. 1).

**Figure 1:**
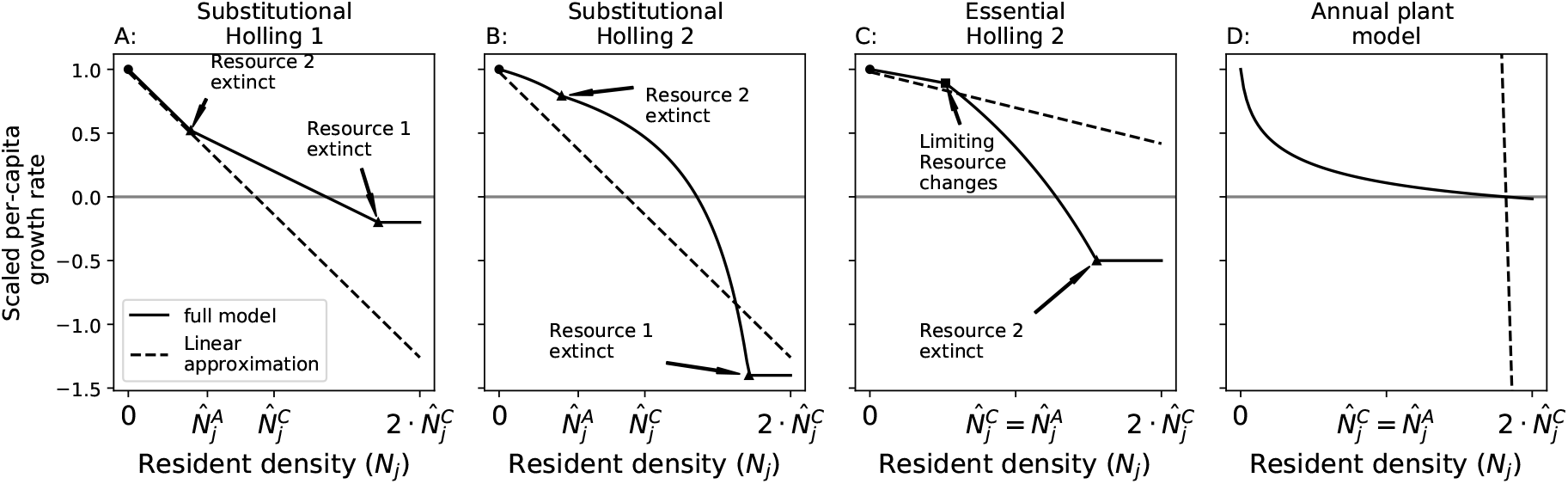
The method to compute 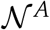 and 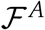 implicitly approximates the actual community model *f_i_*(*N_i_,N_j_*) with a linear per-capita growth rate *r_i_*(1 — *α_ii_N_j_* — *α_ij_N_j_*). We show this approximation explicitly as the scaled per capita growth rates of the focal species (y-axis) as a function of the resident species density (x-axis) for the four full models (solid line = *f_i_* (0,*N_j_*)/*f_i_* (0,0)) and their linear approximations (dashed = 1 — *α_ij_N_j_*). The linear approximations do not capture the full complexity of the models and deviate substantially from the full model. Two main differences between the approximation and the full model stand out: First (A and B), the approximated equilibrium density of the resident 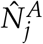 is smaller than the exact resident’s equilibrium density 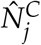 (indicated by different labels for 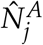 and 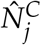). Second (A,B and C), the density for zero-net growth is not equal for the approximation and the full model (intersection with grey line). These two differences occur because the linearisation does not account for resources that go extinct (black triangles, A,B,C) or changes in which essential resource is limiting (black square, C). The full model eventually transforms into a horizontal line (A,B,C) when all resources are extinct and growth rates reduce to the density-independent mortality rates. However, species *j* would not naturally reach densities this high (i.e. above 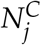). These differences can lead to wrong predictions about the outcome of competition (see Table 2). Chosen parameters (A,B,C) are equivalent to Fig. 2.

When species drive resources to extinction the approximation given in table 1 may lead to an incorrect prediction of coexistence. That is two species may have 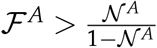 but coexist none the less, or they may have 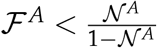 but not coexist. However, in this case the approximation would also lead to different equilibrium densities, i.e. 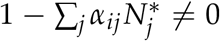, we therefore do not focus further on these cases. However, we caution that one must verify that the linear approximation indeed holds for the equilibrium condition (Appendix 1 and Appendix 2).

### Accounting for complexity enhances understanding of coexistence

Both the approximation and the full niche and fitness differences predict the case where the two species don’t interact and must have 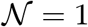 in all four community models. For the other cases 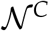 and 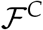 improve our understanding of coexistence.

For certain resource supply rates both species will drive one resource to extinction in mono-culture (Fig. 2 above blue dotted line or below green dotted line). In this case, the ZNGI of the species is non-linear and a linear representation will necessarily loose precision, irrespective of how exactly the linear representation of the model is chosen. When one resource is driven to extinction, both species effectively compete only for one resource and must have have identical feedback loops and no niche differences. For all resource explicit models 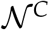 can identify this case (i.e. 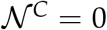). However, 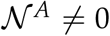 for these resource supply rates for the substitutional resource models (Fig. 2 A and B). Additionally, for the competition for essential resources, the linearisation method predicts 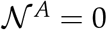 for a community with priority effects (Appendix S2), which contradicts prior findings (Ke & Letten, 2018; Mordecai, 2011; Spaak & De Laender, 2020). Both the approximation and the full 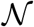 and 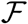 give exactly the same predictions for the absence of niche difference in the annual plant model, i.e. 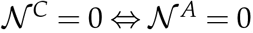, which we interpret as both methods correctly predict if the two species occupy the same niche.

**Figure 2:**
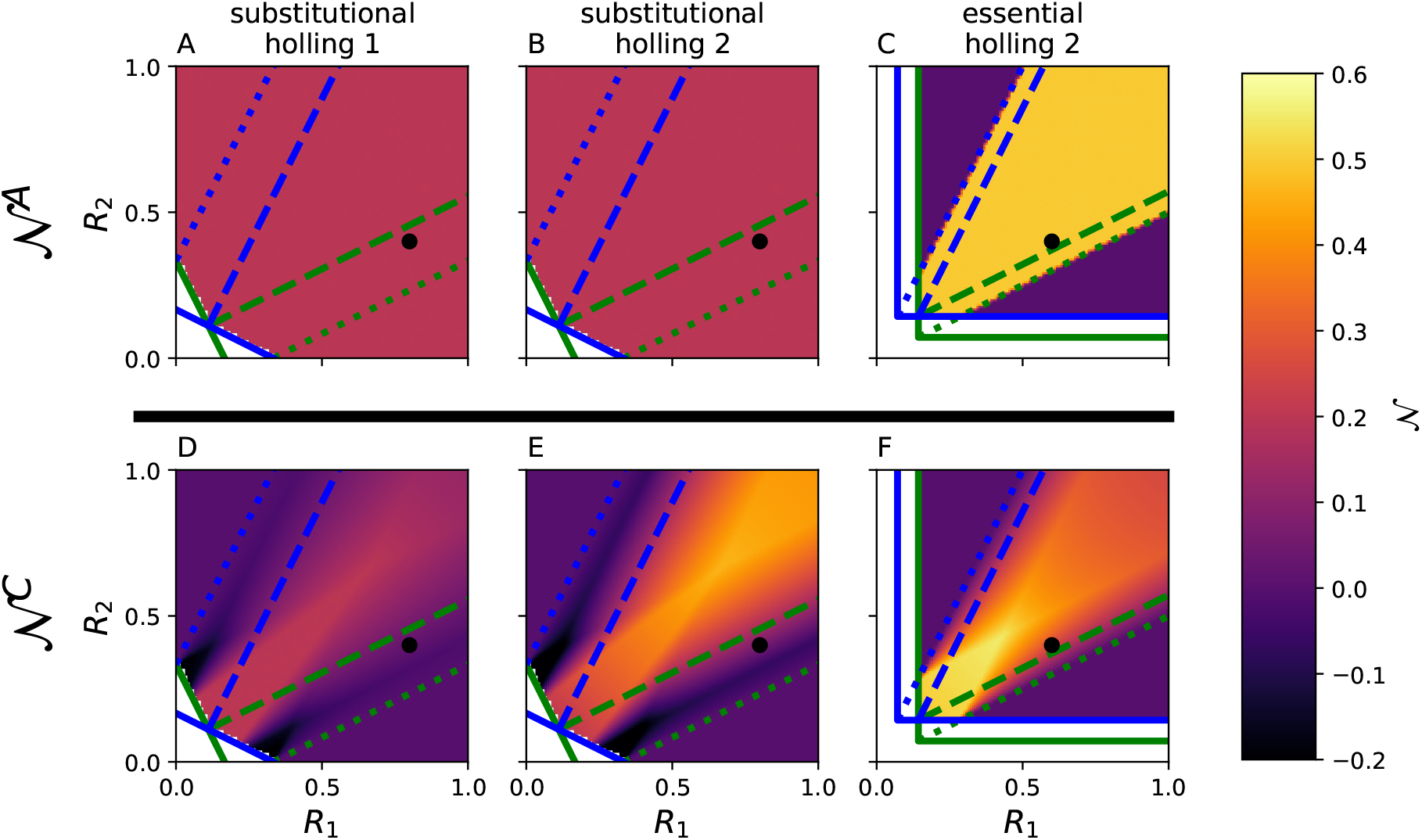
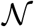 (color) as a function of the resource supply rates for the three different resource explicit community models. 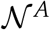 and 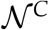 are not defined when one species cannot survive in monoculture (white regions). A,B: 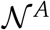 is independent of resource supply rates for substitutable resources. Moreover, the functional response to resource concentrations does not affect 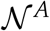 (given *u_ii_* and *w_il_*). In the region where both species are limited by only one resource (above blue dotted and below green dotted line) 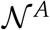 is non-zero, contrary to the assumption that species competing for one resource must have 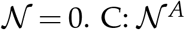 is only affected by changes in the resource supply rates if the limiting resource changes for a species (crossing a dotted line). This change from 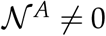 to 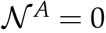, however, is discontinuous. D,E,F: 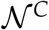 continuously depends on changes in resource supply rates and features local maxima (D-F) and minima (D,E). 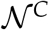 identifies cases where 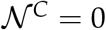 in the region where both species are limited by the same resource. Green and blue solid lines are the ZNGI, dashed lines delimit the coexistence region and dotted lines delimit the region where both species are limited by the same resource. Black dot represents the resource supply rates taken for Fig. 1 and 3.

Any mechanism that is purely equalizing cannot alter the outcome of competition from coexistence to priority effects or vice versa (Ke & Letten, 2018). We found that changes in mortality or resource supply rates can alter the outcome of competition from priority effects to coexistence in certain conditions, and therefore must be stabilizing (Appendix S1). 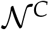 and 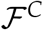 identifies these mechanisms, as they affect 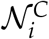 (Fig 3). Conversely, the linear approximations classify any mechanism in the resource competition models that does not affect resource utilisation *u_li_* or the conversion efficiency *w_il_* as purely equalizing; Mechanisms that affect *u_li_* or *w_il_* are both equalizing and stabilizing.

**Figure 3:**
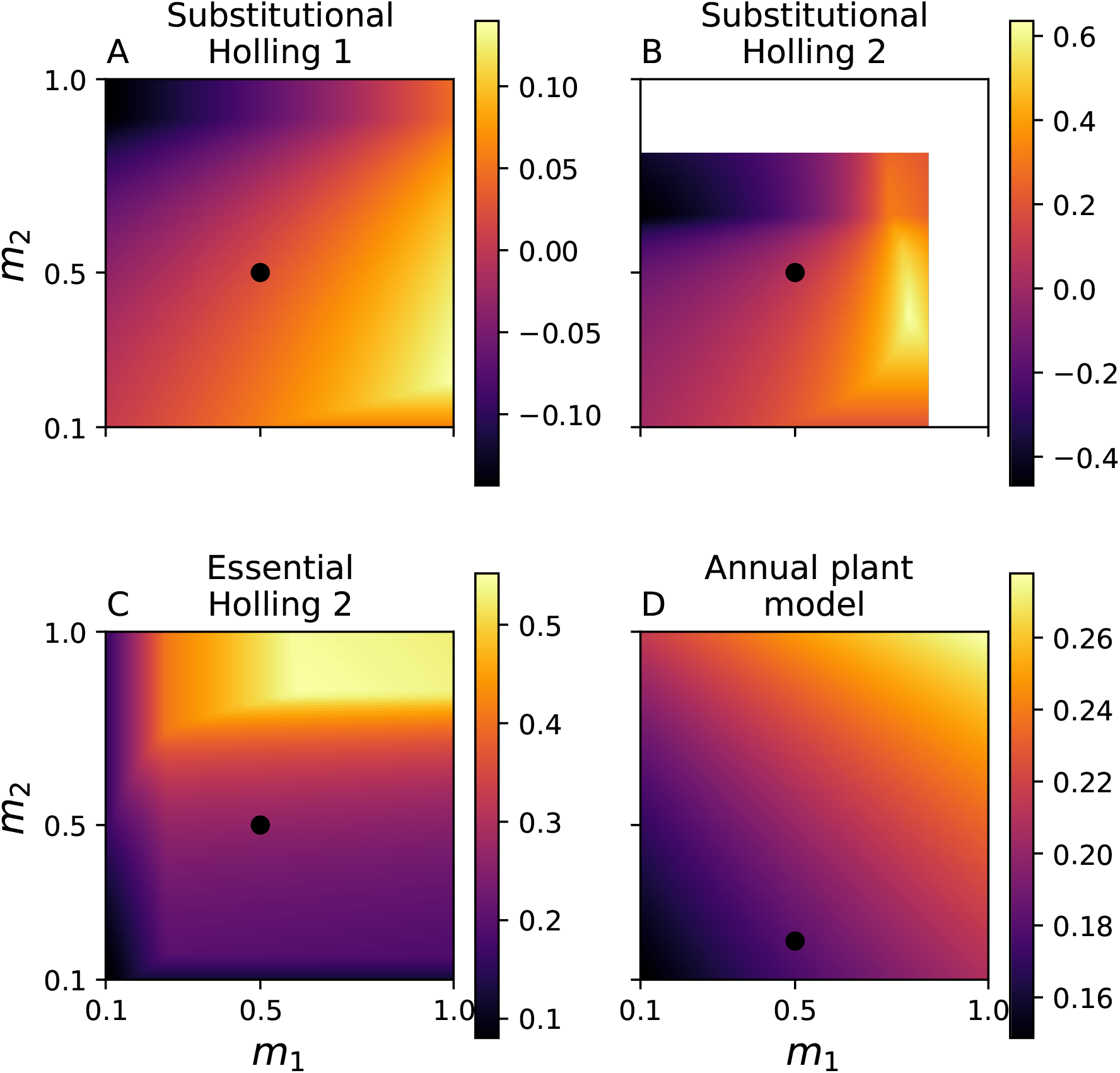
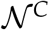 (color) as a function of mortality for the four investigated models. Mortality has a non-monotonic effect on 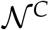 and may lead to local maxima and minima of 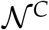 (panel B), similar to changes in the resource supply rates. Blank areas (Panel B) indicate too high mortality for species persistence. Parameters (except mortality) are chosen such as in fig. 2, the black dot represents the same mortality as in fig. 2.

Alternatively, we can see that changes in resource supply rates can be stabilizing in the following scenario: Given two species, the resource supply rates determine whether both species are limited by the same resources (and consequently must have 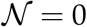) or whether the species coexist (and consequently must have 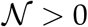). Therefore, changes in resource supply rates can be stabilizing mechanisms. In general, 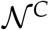 and 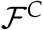 predict that essentially all mechanisms are both stabilizing and equalizing (Song *et al*., 2019).

Importantly, the new interpretation that changes of mortality or resources supply rates are stabilizing, are not based on the specificities of 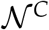 and 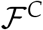. Rather, we draw this argument based on intuitive knowledge of niche differences only. It must therefore apply to any niche and fitness differences definition (Appendix S1).

For the annual plant model interpretation of stabilizing and equalizing mechanisms depends on the method to assess niche and fitness differences. 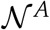 and 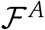 interpret that only mechanisms affecting *α_ij_* can be stabilizing, all other mechanisms are purely equalizing. Conversely, 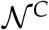 and 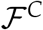 interpret that essentially all mechanisms are both, stabilizing and equalizing, including changes in intrinsic growth rates, seed survival and seed mortality rates (Fig. 3). However, we do not know which of these two interpretations is more useful, as the arguments used for the resource explicit models do not apply for the annual plant model.

### Accounting for complexity reveals new phenomena

A first new insight is the existence of local maxima and minima for 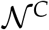. Importantly, the location of these local maxima and minima can be constructed geometrically, similar to how the coexistence region can be constructed (Appendix S3). The existence of these maxima and minima is unexpected, but not counter-intuitive. Imagine the scenario where we keep all community parameters fixed except *S*_1_. For *S*_1_ much smaller than *S*_2_ both species are limited by *R*_1_ only, hence we must have 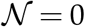, when *S*_1_ and *S*_2_ are comparable the species coexist, hence 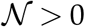, for much larger of *S*_1_ than *S*_2_ the species are limited by *R*_2_, hence 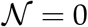. Consequently, 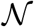 depends in a non-monotonic way on *S*_1_ and will therefore exhibit a maximum. We should therefore expect that 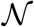 depends in a non-monotonic way on both *S_l_*, as it indeed does.

Another new insight from 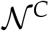 and 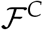 is that competition for substitutable resources can lead to negative 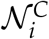, i.e. positive frequency dependence, for certain resource supply rates. However, even for these resource supply rates, the community will not be driven by priority effects. This highlights that negative niche differences (and thus positive frequency dependence) are necessary but not sufficient conditions for priority effects. If fitness differences are too strong, one species will exclude the other, regardless of the initial condition, which is the case in this scenario (Ke & Letten, 2018).

Positive frequency dependence arises when species consume more of the resource that limits their competitor most (Ke & Letten, 2018; Tilman, 1982). For the resource supply rates chosen in figure 2D the blue species drives resource 2 to extinction in monoculture. The green species, as an invader, will therefore only consume resource 1 and have a horizontal utilisation vector. The green species therefore consumes only the favoured resource of its competitor (blue species), which leads to positive frequency dependence (Ke & Letten, 2018). A condition for 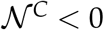 is therefore that resources go extinct, which is why 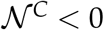 does not occur for essential resources, as species in monoculture can’t drive essential resources to extinction.

## Discussion

An often used approach to compute niche and fitness differences is to represent the equilibrium dynamics of the community model with a linear model (Godwin *et al*., 2020; Godoy & Levine, 2014; Letten *et al*., 2017; Chesson, 2013; Johnson, 2021). We have found that including non-linear species interactions improves our ability to understand coexistence, and yields three important new insights compared to linearisation-based approaches.

First, changes in mortality rates can affect both niche and fitness differences, contrary to earlier results (Chesson, 2000; Godoy *et al*., 2014; Barabás *et al*., 2018; Petry *et al*., 2018; Letten *et al*., 2017). This confirms the earlier findings from Song *et al*. (2019) who found that niche and fitness differences are interdependent. Except in some special cases, changing any parameter always affected both niche and fitness differences. We found no mechanism that acts solely equalizing or stabilizing.

Second, niche differences depend non-monotonically on the resource supply rates and environmental conditions. Again, this result is independent of the specific properties of 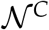 and 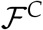, but is based on intuitive properties of niche and fitness differences (see Appendix S1). We also found maxima and minima of niche differences as a function of resource supply. While we were able to show the existence and specific location of such extremes, we know little about their consequences. For example, niche differences are assumed to increase ecosystem function (Carroll *et al*., 2011; Striebel *et al*., 2009) or stability with respect to perturbation (Adler *et al*., 2007). The extent to which these supply rates translate to higher functioning and greater stability remains to be verified.

Third, niche differences based on non-linear community models highlight resource supply rates with negative 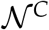 (Fig. 2). At these resource supply rates, species have positive frequency dependence, which was unknown for this well-known community model.

Niche and fitness differences should facilitate understanding of species coexistence by disentangling stabilizing from equalizing mechanisms. We have shown that essentially all mechanisms are both stabilizing and equalizing, as they can alter the outcome of competition from priority effects to coexistence. This is perhaps not surprising when the full non-linear dynamics are taken into account. Nevertheless, we would still anticipate that any given shift of traits or environment conditions will often be more strongly stabilizing or equalizing. It will be valuable in future work to identify the conditions under which equalizing and stabilizing mechanisms exhibit weaker or stronger interdependence. Decomposition methods that account for non-linear interactions will be critical to achieving this objective.

### Future directions of niche and fitness differences

Recent research has shown that non-linear or higher-order species interactions can affect community composition, community stability and coexistence (Grilli *et al*., 2017; Bairey *et al*., 2016; Mayfield & Stouffer, 2017; Singh & Baruah, 2019). We have shown, that niche and fitness differences can be used to understand why non-linear and higher-order species interactions affect community dynamics, however, our results remain limited to simple two-species communities. Applying niche and fitness differences, which include non-linear and higher-order interactions, to more complex multi-species communities is an important next step to understand the effects of non-linear or higher-order effects on population dynamics (Godoy *et al*., 2018; Letten & Stouffer, 2019).

Modern coexistence theory and niche and fitness differences in particular, should predict and understand species coexistence. We have shown that two different methods to asses niche and fitness differences result in different interpretations of the underlying processes. However, there are a total of eleven different methods to assess niche and fitness differences, with little consensus about which to use (Spaak & De Laender, 2020; Godwin *et al*., 2020). Some of these methods can account for non-linear species interactions, some of them cannot. Our results imply that these different methods can potentially lead to different interpretations of the underlying processes. For example, the methods of Carroll *et al*. (2011); Zhao *et al*. (2016) and Carmel *et al*. (2017) interpret essentially all processes as both equalizing and stabilizing, but to different extents (Appendix S4). Similarly, niche differences of Carroll *et al*. (2011) and Zhao *et al*. (2016) depend non-monotonic on resource supply rates, but niche differences of Carmel *et al*. (2017) do not. Additionally, Song & Saavedra (2020) have shown that a mechanism increasing structural niche differences (Saavedra *et al*., 2017) may decrease niche differences as measured by the linearisation approach, that is our interpretation of stabilizing mechanisms depends on the method to assess niche differences.

## Acknowledgements

F.D.L. received support from grants of the University of Namur (FSR Impulsionnel 48454E1) and the Fund for Scientific Research, FNRS (PDR T.0048.16).

## Notes

*Conflict of interest:* The authors declare no conflict of interest.

### Competing Interest Statement

The authors have declared no competing interest.

