## Supplementary material for "The effect of non-linear competitive interactions on quantifying niche and fitness differences"

### Appendix: The effects of linearization on niche and fitness differences

#### Appendix S1: Resource extinction for substitutional resources

Chesson (1990) and Letten *et al.* (2017) have shown, that the linearisation of the community model and the actual community model have the same conditions for coexistence. However, both proofs (implicitly) assume, that none of the resources can go extinct. The resources have a logistic growth term in their community dynamics, it is therefore reasonable to assume, that they are biotic and not abiotic resources and may go extinct, which is probable if we assume multiple resource species (Holt, 1977).

We show, that when we relax this assumption, the linearisation potentially does not correctly predict the outcome of coexistence. The core idea is the same for all counter examples, we will explain the idea in more detail for the case where the linear approximation incorrectly predicts priority effects for a community that coexists. We take two sets of resources (a total of four resources, i.e. $R_1, R_2, R_3$ and $R_4$), for which the two species compete. If the species would only compete for the first set of resources ($R_1$ and $R_2$), they would coexist. Competition only for the second set of resources ($R_3$ and $R_4$) leads to priority effects. The supply rates of the resources of the first set are chosen much higher. At equilibrium, the species will therefore only compete for the first set, and coexist. Conversely, the linearisation approach assumes that the species will compete for all resources, independent of species densities and will therefore incorrectly predict priority effects, assuming parameters were chosen correctly. Figure 1 shows examples where the linear approximation does not correctly predict the outcome of competition.

To see how mortality can affect coexistence, we again take the same set of resources. When mortality is high, then species densities are low and they will not exclude the resources of the second set. As the species compete for all

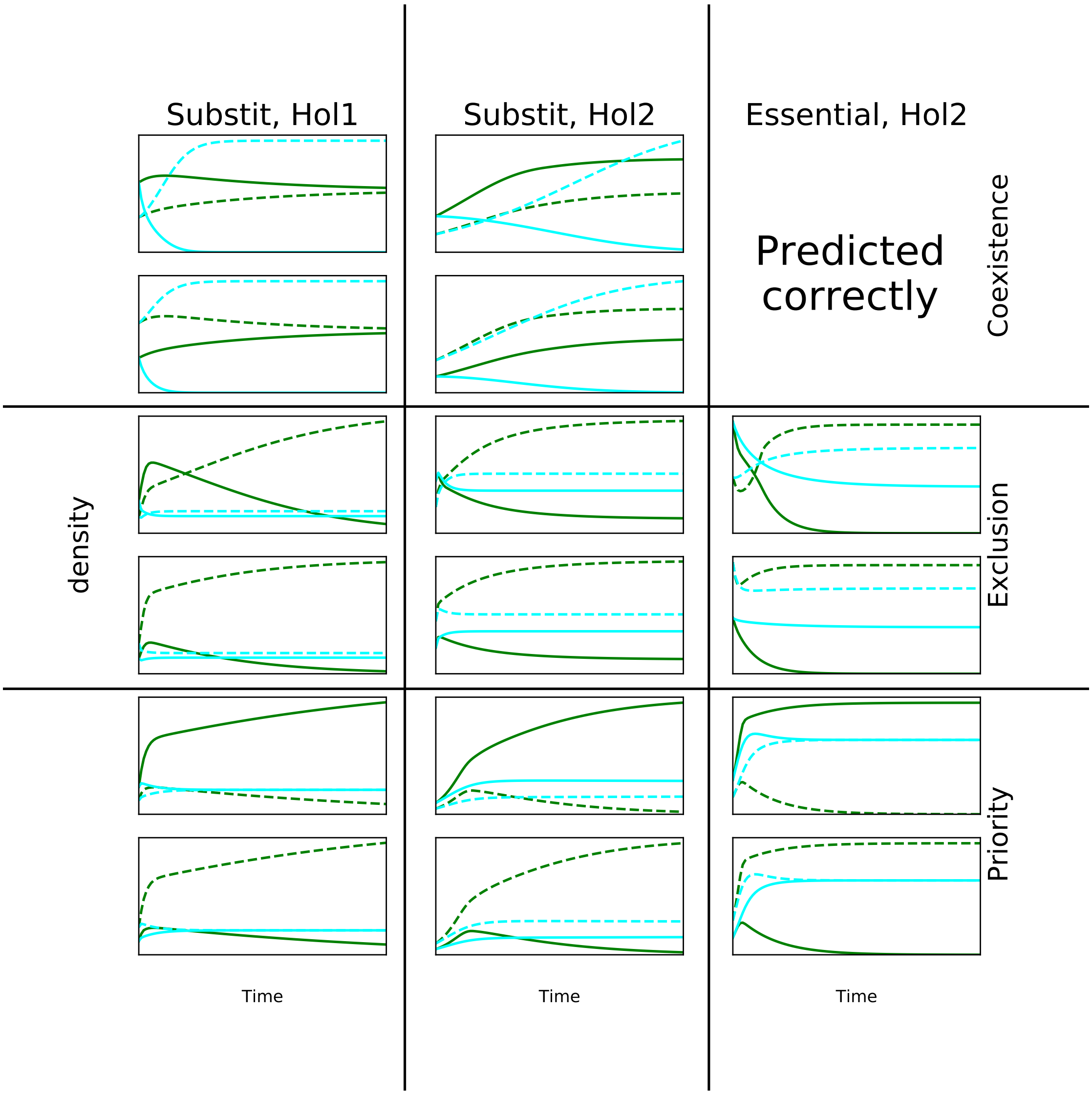


**Figure 1:** For each model (colums) and each outcome of competition (rows) we can find parameter settings in which the linear approximation (red) leads to different predictions about coexistence than the full model (blue). The only exception is species competing for essential resources, in which case coexistence is always predicted correctly. However, the linearisation method of this model may predict coexistence for a community with competitive exclusion or priority effects. The linearisation does not correctly predict the outcome of competition, because resources may go extinct. Shown are densities of species one (full line) and species two (dashed) for the full model (green) and the linear approximation (cyan). The two subplots correspond to different starting densities, to distinguish competitive exclusion from priority effects.

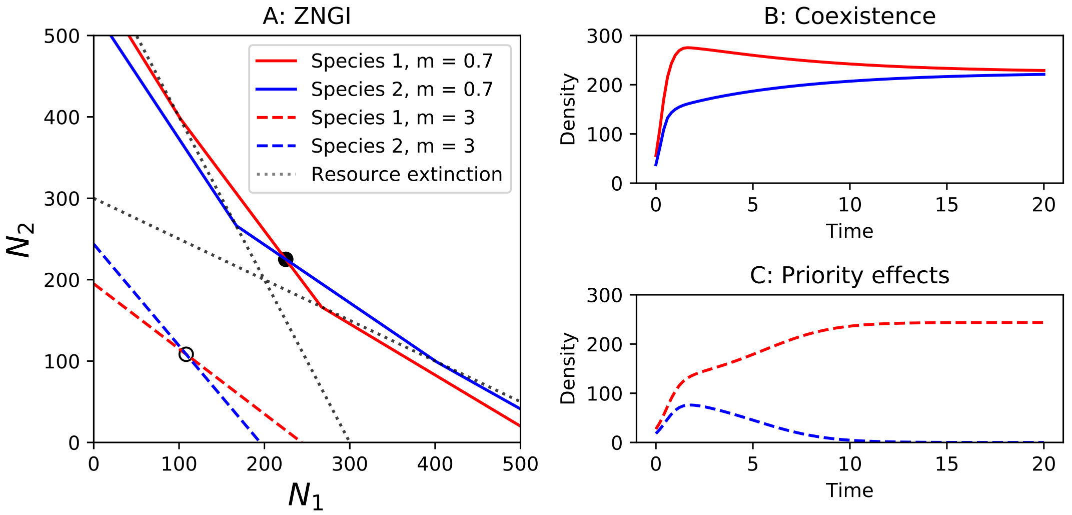


**Figure 2:** A: The zero net growth isoclines (ZNGI) for the two species are shown for different mortality rates. Please note, the x and y-axis show densities of the species one and two, not as in the main text the resource densities. The species compete for a total of 4 resources, a visualisation of the ZNGI as a funciton of the resources is therefore not possible (4 dimensional plot). At low mortality (full lines) species reach higher equilibrium densities and two resources go extinct, as the stable equilibrium is above the resource-extinction line. The two species can stably coexist with the remaining two resources. At high mortality (dashed lines) no resources go extinct and the species comete for all resources, leading to priority effects and an unstable equilibrium. B,C: The species densities over time. Species can coexist under low mortality (B), but not under high mortality (C). The two scenarios differ only in their mortality, but as the outcome of coexistence changes from priority effects to coexistence, mortality must affect niche differences.

resources the winner of competition depends on initial densities (priority effects). Priority effects only occur with negative niche differences (Ke & Letten, 2018; Mordecai, 2011). If we reduce mortality, species densities increase and they exclude the resources of the second set and can coexist. Coexistence, however, is only possible with negative niche differences, consequently mortality can affect niche differences. Similarly, we see that changes in resource supply rates can achieve the same change in competitive outcome.

With a similar idea we create a model with multiple stable equilibria, using six resources, that go extinct under different conditions (Fig. 3). The species coexist, when they compete for resource 3 and 4. Species one can exclude species two, when they compete for resources 1-4 and species two can exclude species one, when they compete for resources 3-6. Additionally, species one has high consumption rate of the resources 5 and 6, conversely, species two has high consumption rates for resources 1 and 2. When both species are at high density, resources 1,2,5 and 6 are extinct and consequently, the species coexist. When only species one is at high density resources 5 and 6 go extinct and species one excludes species two. When only species two is at high density resources 1 and 2 go extinct and species two excludes species one.

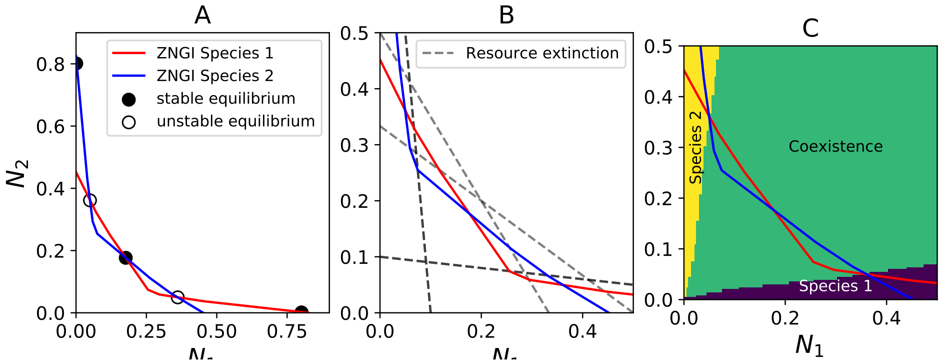


**Figure 3:** A: Species competing for 6 resources feature 5 equilibria. Two unstable (empty circle) and three stable. One equilibrium where species coexist (intersection of ZNGI) and each species in monoculture. B: The multiple equilibria can be explained by the different resources. At each equilibria a different set of resources is extinct, such that the species interactions differ. C: Depending on the starting conditions the community will end up in different equilibrium state, where the species either coexist of exclude each other. The invasion growth rates correctly predict the equilibria at the boundaries, not however that the species can coexist.

### Appendix S2: Essential resources

Letten *et al.* (2017) have shown, that the linearisation of the community model and the actual community model have the same conditions for coexistence. However, both this proof implicitly assumes, that a species is always limited by the same resource independent of species densities. The relaxation of this assumption leads to incorrect predictions about the outcome of competition. There are three qualitatively different ways in which the species can compete for essential resources (Fig. 4). First, the species may coexist or competitively exclude each other, depending on the resource supply ratios (A). In this case the limiting resource is the same at invasion as when no species are present (intrinsic growth rate), then coexistence is predicted correctly (D, proof see 5). Second, the species may have priority effects or exclude each other, depending on the resource supply rations (B). In this case the limiting resource can switch, which leads to wrong predictions about coexistence (E). Third, species one excludes the other species, independent of resource supply ratios (C). Again, in this case the limiting resource can switch and coexistence is not predicted correctly (F).

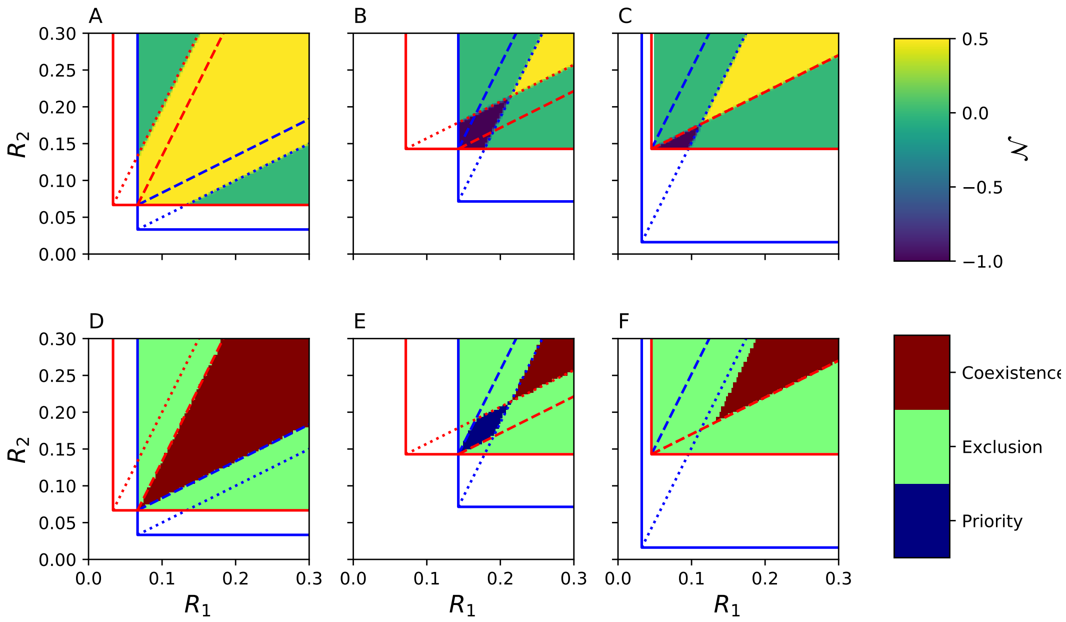


**Figure 4:** $\mathcal{N}^A$ and predictions for competitive outcomes for the three qualitatively different cases of competition for essential resources. The upper row reports $\mathcal{N}^A$, the lower row reports the outcome of competition as predicted by the linearisation approach. A,D: In the first case species coexist when the resource supply ratio is within the dashed lines, which is correctly predicted by the linearisation. B,E: In the second case species either competitively exclude each other (resource supply outside of the dashed lines) or have priority effects. However, the linearisation approach does not correctly predict most of the priority effects and even predicts coexistence and positive niche differences. C,F: In the third case the blue species always excludes the red species, independent of the resource supply ratios. However, the linearisation approach predicts coexistence for some of the resource supply ratios.

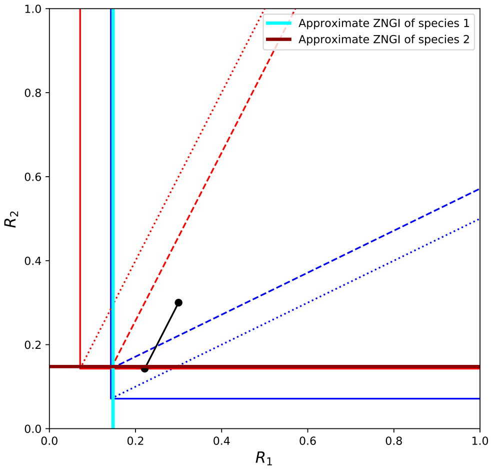


**Figure 5:** The linear approximation leads to a linear ZNGI (dark red and cyan), which approximate the real ZNGI (red and blue). The approximated ZNGI assume that the non-limiting resource is never limiting. Which of the two branches from the correct ZNGI are chosen depends on the resource supply rate, more specifically whether the resource supply rate (black dot) is left or right from the dotted blue and red lines. If the species coexist in the real community (resource supply between dashed lines), then the ZNGI are approximated as shown. These ZNGI lead to coexistence, if and only if the resource supply is between the dashed lines, i.e. the linearisation correctly predicts coexistence.

### Appendix S3: Maxima and minima of niche differences

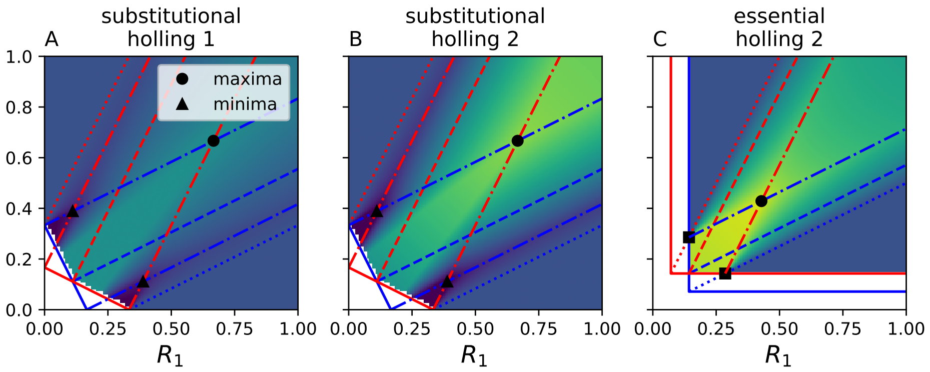


**Figure 6:** The maxima and minima of $\mathcal{N}^C$ can be found geometrically by intersecting resource consumption vectors. A,B,C: In the top right region (delimited by the dash dotted lines intersection at the black dot) the invasion growth rates of both species are maximal and constant. Of all the resource supply points within this region the intrinsic growth rates are minimal for both species at the black dot. As niche differences increase with invasion growth rates and decrease with the intrinsic growth rates this is the location where niche differences are maximal. A,B: Competition for substitutional resources features two local minima (one of which is global) and one global maximum. The dash-dotted lines are parallel to the resource consumption vectors of the corresponding colour and intersect the resource axis at the ZNGI. The intersection of two such lines lead to a local maxima (black dot) or minima (black triangle). The minimal of niche differences are located where one species has maximal and the other species has a minimal invasion growth rate. C: Again, the global maxima of $\mathcal{N}^C$ is located at the intersection of two resource consumption vectors (dash-dotted lines). These resource consumption vectors are anchored where the dotted consumption vectors intersect the ZNGI of the other species (black square).

### Appendix S4: Comparing methods to compute niche and fitness differences

There are other methods to compute niche and fitness differences. The most often used definition not investigated in the main text is by Carroll *et al.* (2011). Their method can be understood as a linear approximation of the community model, with $\frac{\alpha_{ij}}{\alpha_{jj}} = \frac{f_i(0,N_j^*)}{f_i(0,0)}$, where $f_i(0,N_j^*)$ is the invasion growth rate and $f_i(0,0)$ is the intrinsic growth rate. Two other methods use the invasion growth rates to define niche and fitness differences. All these methods are based on invasion growth rates, therefore they correctly predict the outcome of competition in the cases analysed, similarly to the method based on the full model. However, all of these methods have non-zero niche differences, when the two species compete for only one resource.

Again other definitions are based specific community models, notably linear community models (see Spaak & De Laender (2020) for a review). We do not review these, as they all require to first fit a linear model which comes with the disadvantages discussed in the main text.

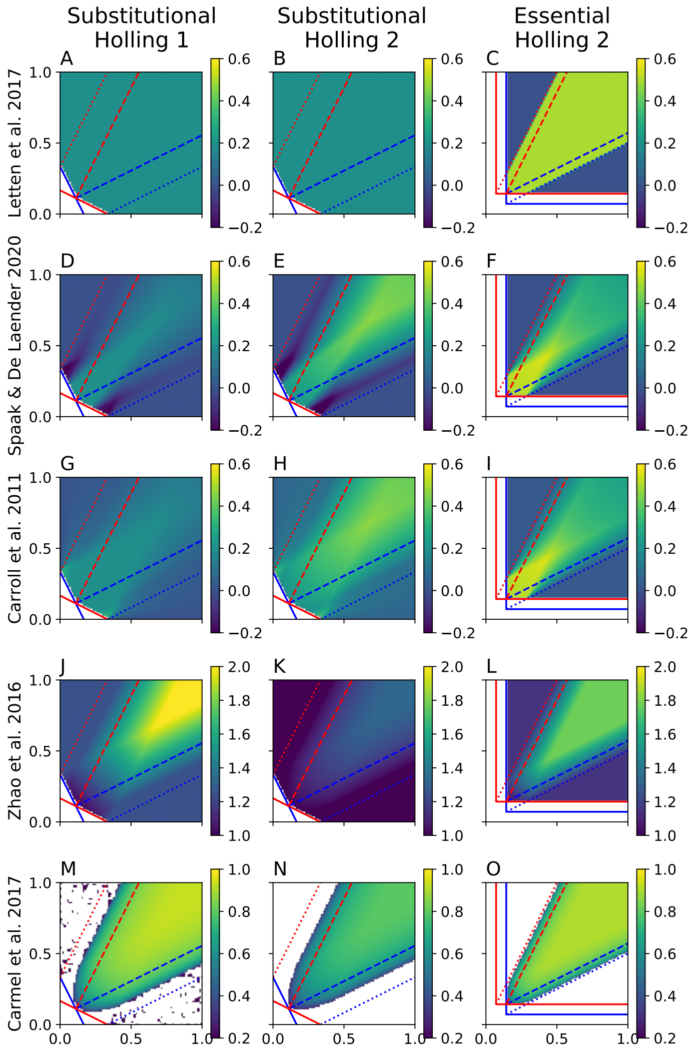


**Figure 7:** Niche differences for five different methods (rows) and 3 different models (colums). Spaak & De Laender (2020) is the only method that predicts zero niche differences when species compete only for one resource (outside the dotted lines). All methods except the linearisation methods interpret changes in resource supply rates as stabilizing, that is they are sensitive to at least certain non-linear dynamics. The definition of Carmel *et al.* (2017) is not defined for some cases where the species can survive in monoculture (white region). Please note that the color scales differ per method.

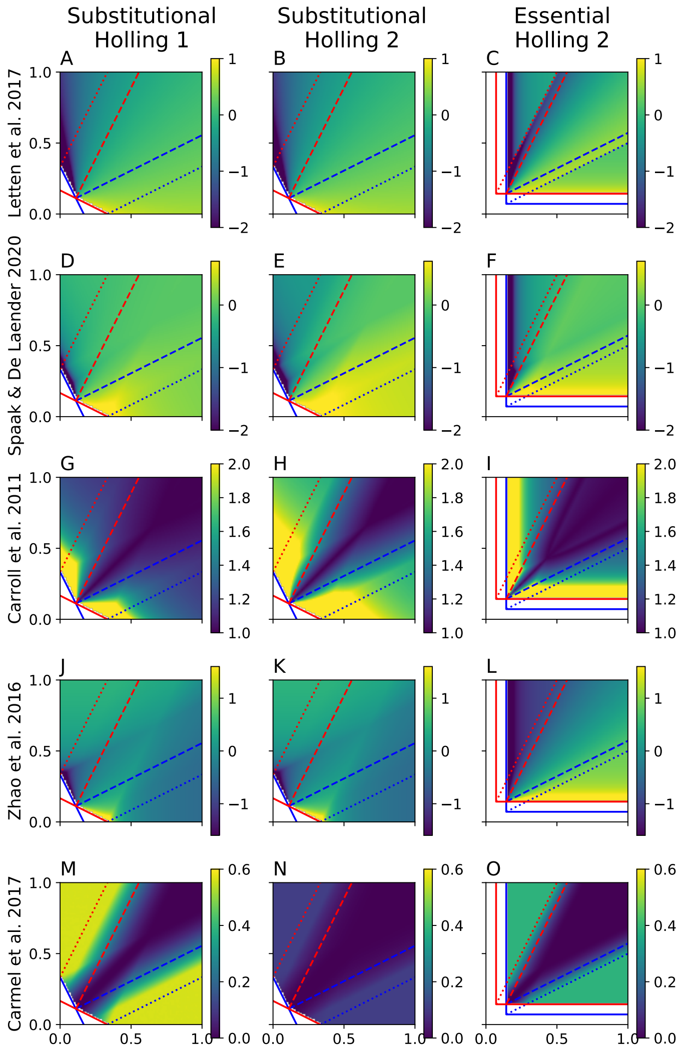


**Figure 8:** Fitness differences for five different methods (rows) and 3 different models (colums). The definitions also differ for the interpretation of competitive equivalence. For the definitions Letten *et al.* (2017); Spaak & De Laender (2020); Zhao *et al.* (2016) and Carmel *et al.* (2017) species with equal competitive strength with have zero fitness differences, while for Carroll *et al.* (2011) fitness differences of one imply competitive equality. Please note that the colorscales differ per method.

### Appendix S5: The annual plant model

Godoy & Levine (2014) show that niche and fitness differences as defined by them correctly predict coexistence. We show here, that the two methods to compute niche differences agree on when the two species occupy the same niche. Assume $\mathcal{N}^A = 0 \Leftrightarrow \sqrt{\frac{\alpha_{ij}\alpha_{ji}}{\alpha_{ii}\alpha_{jj}}} = 1$. We can then choose $c_j = \frac{g_i}{g_j}\sqrt{\frac{\alpha_{jj}\alpha_{ij}}{\alpha_{ji}\alpha_{ii}}}$ which results in:

$$\begin{aligned}
\exp(f_i(0,N_j^*)) &= (1-g_i)s_i + \frac{\lambda_i g_i}{1+\alpha_{ij}g_jN_j^*} && (1)\\
&= (1-g_i)s_i + \frac{\lambda_i g_i}{1+\sqrt{\frac{\alpha_{jj}\alpha_{ii}}{\alpha_{ji}\alpha_{ij}}}\alpha_{ij}g_jN_j^*} && (2)\\
&= (1-g_i)s_i + \frac{\lambda_i g_i}{1+\alpha_{ii}\sqrt{\frac{\alpha_{jj}\alpha_{ij}}{\alpha_{ji}\alpha_{ii}}}g_jN_j^*} && (3)\\
&= (1-g_i)s_i + \frac{\lambda_i g_i}{1+\alpha_{ii}g_ic_jN_j^*} && (4)\\
&= \exp(f_i(c_jN_j^*,0)) \Rightarrow \mathcal{N}^C = 0 && (5)
\end{aligned}$$

Conversely, assume that $\mathcal{N}^C = 0$, then we have $f_i(0,N_j^*) = f_i(c_jN_j^*,0) \Leftrightarrow \alpha_{ij}g_jN_j^* = \alpha_{ii}g_ic_jN_j^*$ and similarly $\alpha_{ji}g_iN_i^* = \alpha_{jj}g_jc_iN_i^*$, which leads together with the equation $c_j = c_i^{-1}$ to $\sqrt{\frac{\alpha_{ij}\alpha_{ji}}{\alpha_{ii}\alpha_{jj}}} = 1$.

### Appendix S6: How to compute $\mathcal{N}^C$ and $\mathcal{F}^C$ in resource explicit models

Spaak & De Laender (2020) define niche and fitness differences for a community in which the species growth rates depend directly on the species densities, i.e. $\frac{1}{N_i}\frac{dN_i}{dt} = f_i(N_i, N_j)$. In the resource explicit models the species growth rates do not depend directly on the species densities, but rather on the resources densities and therefore indirectly on the species densities. To compute $\mathcal{N}^C$ and $\mathcal{F}^C$ we have to compute the intrinsic growth rate ($f_i(0,0)$), the invasion growth rate ($f_i(N_j,0)$) and the no-niche growth rate ($f_i(c_jN_j,0)$). The invasion growth rate of the species is computed, when the resident species $j$ is at equilibrium, this implies that the resources are also at equilibrium. Similarly, the intrinsic growth rate is the growth rate when no species has yet consumed any resources, consequently all resources are at carrying capacity (and also at equilibrium). Finally, the no-niche growth rate is the growth rate, when species $i$ is at the converted equilibrium density of species $j$, therefore the resources are again at equilibrium. That is, to compute $\mathcal{N}^C$ and $\mathcal{F}^C$ we can assume that the resources are at equilibrium. However, we do not assume that the dynamics of the resources are faster than the dynamics of species (time-scale separation), rather all growth rates are evaluated at equilibrium.

We illustrate how to compute $\mathcal{N}^C$ and $\mathcal{F}^C$ for two species competing for substitutable resources with Holling type 2 response functions. First we set the resource densities to equilibrium, i.e. $\frac{dR_l}{dt} = 0 = S_l - R_l - \sum_j u_{lj}N_j$, which leads to $R_l = S_l - \sum_j u_{lj}N_j$. Note this equation holds only for all resources that are not extinct, all other resources have $R_l = 0$, for notational convenience we assume that no resources go extinct. We now compute the equilibrium density of species $j$:

$$\frac{1}{N_j} dN_j dt = 0 = \frac{\sum_l w_{jl} R_l}{k_j + \sum_l w_{jl} R_l} - m_j \tag{6}$$

$$= \frac{\sum_l w_{jl} \left(S_l - u_{lj} N_j\right)}{k_j + \sum_l w_{jl} \left(S_l - u_{lj} N_j\right)} - m_j \tag{7}$$

$$m_j \left( k_j + \sum_l w_{jl} \left(S_l - u_{lj} N_j\right) \right) = \sum_l w_{jl} \left(S_l - u_{lj} N_j\right) \tag{8}$$

$$m_j k_j + m_j \sum_l w_{jl} S_l - m_j \sum_l w_{jl} u_{lj} N_j = \sum_l w_{jl} S_l - \sum_l w_{jl} u_{lj} N_j \tag{9}$$

$$(1 - m_j) \sum_l w_{jl} u_{lj} N_j = (1 - m_j) \sum_l w_{jl} S_l - m_j k_j \tag{10}$$

$$N_j = \frac{\sum_l w_{jl} S_l - \frac{m_j k_j}{1 - m_j}}{\sum_l w_{jl} u_{lj}} \tag{11}$$

This is of course exactly $\frac{1}{\alpha_{jj}}$ from table 1, as we assumed that no resources go extinct. We can now compute the invasion growth rate of species $i$ with

$$f_i(0, N_j^*) = \frac{\sum_l w_{il} R_l}{k_l + \sum_l w_{il} R_l} - m_i \tag{12}$$

$$= \frac{\sum_l w_{il} \left( S_l - u_{lj} N_j^* \right)}{k_l + \sum_l w_{il} \left( S_l - u_{lj} N_j^* \right)} - m_i = \frac{\sum_l w_{il} \left( S_l - u_{lj} \frac{\sum_l w_{jl} S_l - \frac{m_j k_j}{1 - m_j}}{\sum_l w_{jl} u_{lj}} \right)}{k_l + \sum_l w_{il} \left( S_l - u_{lj} \frac{\sum_l w_{jl} S_l - \frac{m_j k_j}{1 - m_j}}{\sum_l w_{jl} u_{lj}} \right)} - m_i \tag{13}$$

The intrinsic growth rate is given by

$$f_i(0,0) = \frac{\sum_l w_{il} S_l}{k_i + \sum_l w_{il} S_l} - m_i \tag{14}$$

Again, this is identical to $r_i$ from table 1.

Finally, to compute the no-niche growth rate we have to solve the following equation for $c_j$:

$$\frac{\frac{\sum_l w_{il}\left(S_l-u_{lj}N_j^*\right)}{k_l+\sum_l w_{il}\left(S_l-u_{lj}N_j^*\right)}-m_i-\left(\frac{\sum_l w_{il}\left(S_l-u_{lj}c_jN_j^*\right)}{k_l+\sum_l w_{il}\left(S_l-u_{lj}c_jN_j^*\right)}-m_i\right)}{\frac{\sum_l w_{il}(S_l)}{k_l+\sum_l w_{il}(S_l)}-m_i-\left(\frac{\sum_l w_{il}\left(S_l-u_{lj}c_jN_j^*\right)}{k_l+\sum_l w_{il}\left(S_l-u_{lj}c_jN_j^*\right)}-m_i\right)}= \tag{15}$$

$$\frac{\frac{\sum_l w_{jl}\left(S_l-u_{li}N_i^*\right)}{k_l+\sum_l w_{jl}\left(S_l-u_{li}N_i^*\right)}-m_j-\left(\frac{\sum_l w_{jl}\left(S_l-u_{li}c_iN_i^*\right)}{k_l+\sum_l w_{jl}\left(S_l-u_{li}c_iN_i^*\right)}-m_j\right)}{\frac{\sum_l w_{jl}(S_l)}{k_l+\sum_l w_{jl}(S_l)}-m_j-\left(\frac{\sum_l w_{jl}\left(S_l-u_{li}c_iN_i^*\right)}{k_l+\sum_l w_{jl}\left(S_l-u_{li}c_iN_i^*\right)}-m_j\right)} \tag{16}$$

While this equation may look very daunting, it's actually reasonably simple. $c_j = \frac{1}{c_i}$ is the only variable, everything else are parameters. The equation cannot be solved analytically, but very easily with numerical methods.
